# Significant SAR11 removal by a hard-bottom community

**DOI:** 10.1101/2022.12.24.521757

**Authors:** Ayelet Dadon-Pilosof, Keats Conley, Marcelino T. Suzuki

**Affiliations:** The Faculty of Marine Science, Ruppin Academic Center, Michmoret, Israel; Department of Ecology, Evolution & Behavior, The Hebrew University of Jerusalem, Israel; University of Oregon, Oregon Institute of Marine Biology - 1585 E 13th Ave, Eugene, OR, USA 97403; Sorbonne Université, CNRS, Laboratoire de Biodiversité et Biotechnologies Microbiennes, LBBM, Observatoire Océanologique, F-66650, Banyuls-sur-mer, France

## Abstract

Microorganisms are a key component of the marine food webs through the microbial loop. In previous work, we have shown that some bacteria, including Candidatus *Pelagibacter* spp (SAR11)—the most abundant bacterium in the ocean—can evade filtration by benthic and planktonic tunicates. Here we tested whether differential removal of microbial taxa by benthic filter-feeders can be observed in the distribution and abundance of microbial taxa from hard-bottom subtidal communities, a common coastal habitat in the Eastern Mediterranean Sea towards the open sea. The abundance of microbial groups along crosss-hore transects was measured based on combined flow cytometry and SSU rRNA gene metabarcoding. Our results show that most groups were depleted (up to 50%) over the hard-bottom compared to the open sea, but unexpectedly we did not observe a clear differential removal of different taxa, SAR 11 notably. This study indicates a strong top-down control of the abundance of pelagic microorganisms over shallow hard-bottom where suspension feeders are common.

## Introduction

Marine microbial communities form the basis of the ocean food web and mediate most of the energy and material fluxes in the ocean (Glöckner et al. 2012). Microorganisms constitute a large fraction of the living biomass in the sea (Pomeroy et al. 2007), and the structure and function of their populations are shaped by a delicate balance between growth and mortality (Pernthaler 2005). Grazing and virus-driven lysis constitute the main sources of mortality (Sánchez et al. 2020), countered by the capabilities of microorganisms to avoid grazing (Matz and Kjelleberg 2005), survive digestion or resist viral lysis. Grazing or predation on microorganisms by either planktonic or benthic organisms is an important mortality factor in many habitats (e.g., Verity 1991; Gili and Coma 1998; Gorsky et al. 1999; Riisgard and Larsen 2001; Pernthaler 2005; Patten et al. 2011a). In the pelagic realm, protists commonly dominate the guild of grazers of bacteria (Calbet and Landry 2004; Matz and Kjelleberg 2005). However, in some habitats and seasons, grazing by metazoan and lysis by phages may dominate mortality (Hahn and Höfle 2001).

Bacteria form some 65-86% of the biomass of microorganisms in the upper ocean (Morris et al. 2002). However, only a small number of bacterial groups dominate that guild (Teeling et al. 2012). In the oligotrophic eastern Mediterranean, pico-cyanobacteria (*Synechococcus* and *Prochlorococcus*) and a few members of Candidatus *Pelagibacter ubique* (SAR11) clade dominate microbial communities, accounting for >70% of the total bacterial biomass (Partensky et al. 1999, Dadon-Pilosof et al. 2017). SAR11 is a clade of heterotrophic bacteria, which constitutes 15–60% of total bacteria in the upper ocean (Morris et al. 2002,2012; Rappé et al. 2002; Eiler et al. 2009; Giovannoni 2017). It is one of the smallest free-living bacteria in the sea and is thought to be the most abundant group in the world ocean. Within that group, some ecotypes thrive in oligotrophic environments while others in more productive waters (Morris et al. 2002, Salter et al 2015). Members of the SAR11 clade also have the lowest nucleic acid content (LNA) among all non-photosynthetic bacteria (Mary et al. 2006). Dadon-Pilosof et al. (2017) reported that some bacteria, especially members of the SAR11 clade, can effectively evade grazing from both pelagic and benthic tunicates and this lack of grazing pressure on SAR11 could partially explain its abundance and ubiquity.

A diverse guild of benthic invertebrate suspension feeders, including sponges, bivalves, cirripedians cnidarians, bryozoans, and tunicates, often dominates subtidal hard substrates (Topçu et al. 2010). Their diet ranges from consumption of dissolved organic matter (DOM) through grazing on microorganisms such as phytoplankton, virioplankton, archaea and bacteria, as well as feeding on zooplankton and detritus (Gili and Coma 1998; Topçu et al. 2010). Hard-bottom subtidal communities along the Mediterranean Sea are diverse and are undergoing dramatic changes in the recent decades due to the combined effect of global warming, overfishing, and the introduction of invasive species (Rilov et al. 2019). Within these communities, retention of small particles is an adaptive advantage since picoplankton often dominate the planktonic community biomass (Topçu et al 2010).

Grazing by benthic suspension feeders on microorganisms is also an important component of the benthic-pelagic coupling in coral reefs (Yahel et al. 1998; Genin et al. 2002, 2009; Patten et al. 2011). This grazing pressure on the microbial community is not necessarily uniform as it depends on the spatial heterogeneity of the distribution of different suspension feeders and their respective diets (Yahel et al. 2006, 2009; Hanson et al. 2009; Dadon-Pilosof et al. 2017). Differential capture of particles from the ambient water based on their size, concentration or morphological features is therefore expected to be reflected in the prey distribution (Gili and Coma 1998).

The goal of the current study was to indirectly evaluate the effect of the whole benthic community removal on the distribution of microorganisms across a shallow, subtropical rocky coast. Following the study of Yahel et al. (1998) across a coral reef, a study that preceded the demonstration of the role of phytoplankton grazing in the trophic dynamics of coral reefs (Genin et al. 2009; Monismith et al. 2010), we sought to test the hypothesis that differential removal on microbes is also reflected in the cross-shore distribution of bacteria and other picoplankton.

## Methods

### Study site

Sampling was conducted off Michmoret, Israel (32° 24′N, 34° 52′E) in the Eastern Mediterranean Sea. The oligotrophic Israeli shoreline is characterized by extremely low nutrient levels (Nitrate 0.5-1 µM, Phosphate 0.05-0.1 µM, Krom and Suari 2015), high salinity (38.3 to 40.0 PSU) and relatively warm waters (16.5 to 30.8°C) (Suari et al. 2019) that is dominated in terms of numbers and biomass by pico- and nanoplankton sized organisms (Herut et al. 2000, Raveh et al. 2015). Water stratification appears usually in spring following a deep winter mixing that is enhanced by the high salinity of surface waters.

Eight transects were performed between December 2015 and April 2017 (Table S1). Each transect encompassed eight stations spanning from 1.5 m depth at the hard-bottom subtidal towards the “open-sea” (∼ 0.9 km off-shore, water depth >12 m, Fig. 1). During the sampling periods the average temperature (±SD) was 19.7±0.7 °C, salinity was high (39.25±0.2 PSU), and the water column was fully oxygenated (dissolved oxygen concentration 216±6 µM). Due to the need to work very close to the bottom at the rocky shoreline, sampling dates were dictated by sea conditions and were limited to days of calm sea (wave height < 30 cm) and weak winds (<6m s^-1^). To get a larger scale context within the eastern Mediterranean shelf offshore of the high-resolution cross-shore transects, six cross-shelf transects were sampled during March 2018, spanning from one km offshore, equivalent to the farthest station in the small scale transects, to 42 km west to the shelf edge at 750 m bottom depth (Fig S1).

**Figure 1:**
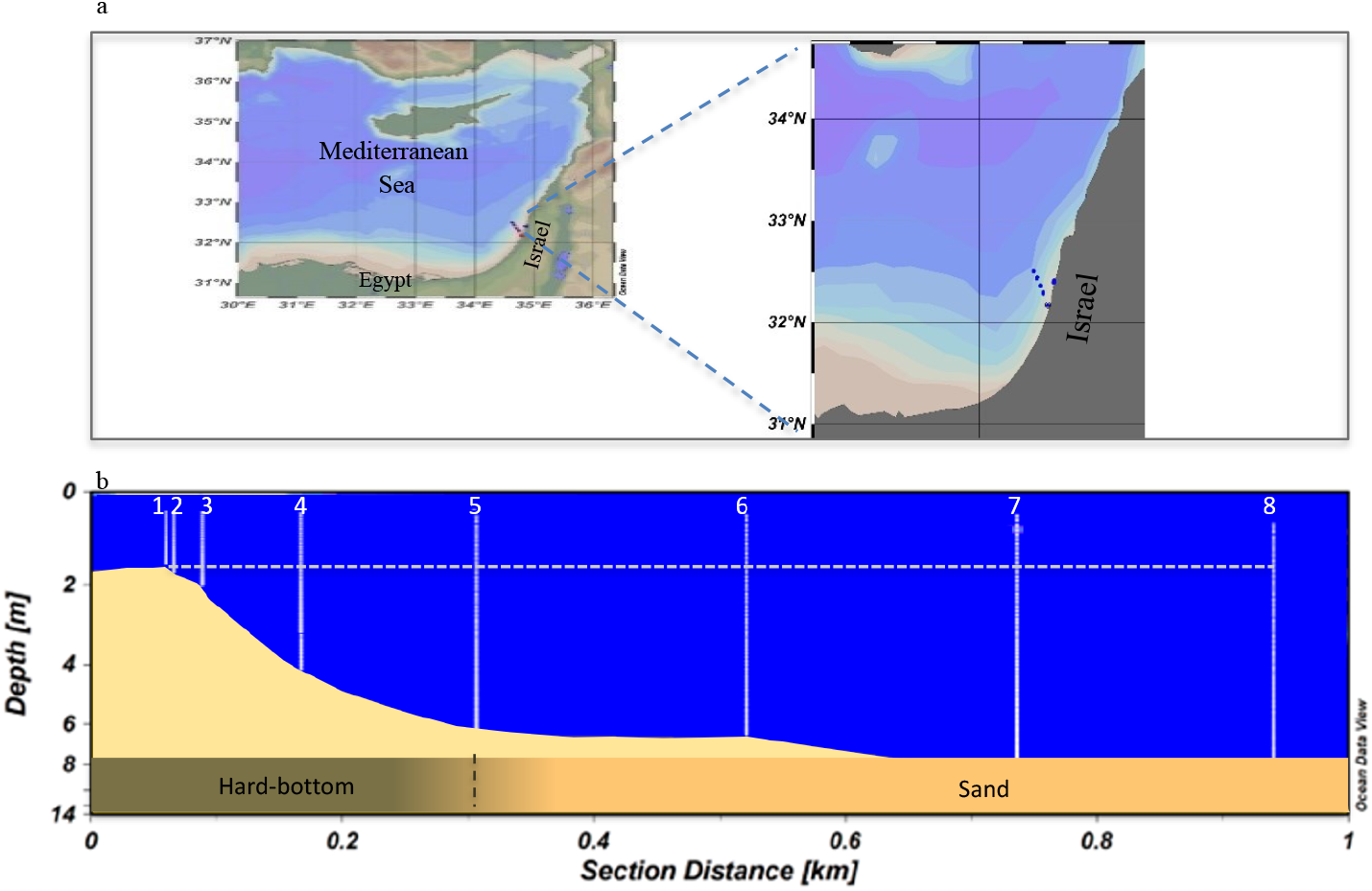
The location of the transect line in the Eastern Mediterranean (a) transects location along the Israeli shore (b) transect showing the bottom (yellow), sampling stations (white numbered lines), dotted white line (1.5 m depth) representing sampling depth.

The shallowest sampling station in each transect was located ∼150 m offshore at 1.6-1.8 m depth where Niskin bottle was placed just above the rocky bottom (c.a. 10 cm) to collect a sample. The 1.5 m sampling depth was retained along the transect as the water deepened down to 12 m at the furthest offshore station, ∼0.9 km offshore. The four shallowest stations (hereafter Stations 1-4) were located above hard bottom (rocky area), the 5^th^ station was located at the boundary between rocks and sand, while stations 6 to 8 were above sandy bottom.

### Sampling methods

Prior to sampling, each station was marked with a moored surface buoy and profiles of temperature, salinity, fluorescence, and oxygen were measured using a Seabird SBE19plusV2 CTD equipped with an *in vivo* chlorophyll fluorimeter (Cyclop7, Turner Designs) and a dissolved oxygen sensor (SBE43, Seabird). The CTD was lowered in a horizontal position at a slow pace (about a meter per minute) so that the water column was adequately profiled even in the shallow-most stations. Due to the need to work close to the bottom at the rocky shoreline, sampling was conducted from either a kayak or a small inflatable skiff.

Seawater was collected simultaneously with CTD profiles, using a 5 L Niskin bottle. At each station, samples were collected directly from the Niskin bottle for all the required analyses: flow cytometry, DNA extraction and Chl-a measurements. Water samples were kept on ice in the dark for further processing in the laboratory within 2-3 hours. Samples for DNA were collected only during three of the eight sampling times.

### Sample analysis

#### Flow cytometry

Flow cytometry was used to quantify the concentrations and the cell characteristics of non-photosynthetic microorganisms (hereafter referred to as non-photosynthetic bacteria), and the following four dominant autotrophic groups: *Prochlorococcus* (Pro), *Synechococcus* (Syn), pico-eukaryotic algae (PicoEuk), and nano-eukaryotic algae (NanoEuk). We used an Attune® Acoustic Focusing Flow Cytometer (Applied Biosystems) equipped with a syringe-based fluidic system that allows a precise adjustment of the injected sample volume and hence high precision of the measurements of cell concentrations (±5%). The instrument’s optical unit contained violet and blue lasers (405 and 488 nm, respectively) and was further adapted for the analysis of marine ultra-plankton samples as described below.

Aliquots of 1.8 mL were collected from each water sample and transferred into 2 mL cryotubes (Corning cat No. 430659). Samples were first incubated for 15 min at room temperature with Glutaraldehyde 50% (electron microscopy grade, Sigma-Aldrich, cat No. 340855) at 0.1% (final concentration). Samples were frozen in liquid nitrogen (at least 60 min) and then stored at -80 °C until analysis (within a few days).

Each sample was analyzed twice. First, 600 µl of the sample water was analyzed at a high flow rate (100 µL min^-1^) for the determination of ultra-phytoplankton with a dual threshold (trigger) on the red fluorescence channels of the violet and blue lasers. A second run was used to analyze cells with no autofluorescence, i.e., non-photosynthetic microbes. To visualize these cells, a 300 µL aliquot of the sampled water was incubated with the nucleic acid stain SYBR Green I (20-120 min dark incubation at room temperature, 1:10^4^ of the SYBR Green commercial stock). For this run, we used a low flow rate of 25 µL min^-1^ and the instrument was set to high sensitivity mode. Seventy-five µL of the sample water was analyzed with a dual threshold (trigger) on green fluorescence channels of the violet and blue lasers. The taxonomic identification was based on orange fluorescence (Bl2, 574±13 nm) of phycoerythrin and red fluorescence (Bl3, 690±20 nm and VL3, 685±20 nm) of chlorophylls; side-scatter (SSC), provided a proxy of cell surface complexity and cell volume (Marie et al. 1999), and forward-scatter (FSC) was a proxy of cell size (Cunningham and Buonnacorsi 1992; Simon et al. 1994).

Where possible, the non-photosynthetic bacteria were further divided based on their green fluorescence (proxy for nucleic acid content) and forward scatter (proxy for size) into three groups: LNA, low nucleic acid non-photosynthetic bacteria; HNA-Ls, high nucleic acid low-scatter non-photosynthetic bacteria; HNA-Hs, high nucleic acid high-scatter non-photosynthetic bacteria (Zubkov et al. 2004). Similarly, the eukaryotic algae were separated to pico- and nano-phytoplankton (Simon et al. 1994). The size of *Synechococcus* is still somewhat controversial, indicating a range of 0.3 to 1.2 µm (e.g., Uysal 2001; Garcia et al. 2016). For pico-eukaryotic algae, we followed Worden and Not (2008), who suggested a size range of up to 3.0 µm. Larger cells were termed nano-eukaryotic algae (2.0-20 µm). As a rough proxy of cell size, we used the ratio of the median forward scatter of each cell population to that of the median forward scatter of reference beads (Polysciences™, cat# 23517, Flow Check High-Intensity Green Alignment 1.0 µm) that were used as an internal standard in each sample. See Dadon-Pilosof et al. (2019) for a further discussion of the accuracy of size estimates.

### Chlorophyll measurements

Water samples (∼300 mL) were collected directly from the Niskin bottles at each station into dark volumetric BOD glass bottles (Wheaton 227667) and maintained on ice in dark cool box. In the lab, samples were prefiltered through a 100 μm mesh (to remove large zooplankton and/or aggregates and suspended pieces of benthic algae) and filtered using low vacuum onto a 25 mm Whatman GF/F filter. Filters were kept frozen at -20°C in 20 mL scintillation vials until further processing. To ensure complete chlorophyll *a* extraction of coastal phytoplankton, we used a hot dimethyl sulfoxide (DMSO) extraction method (Burnison 1980). Briefly, 2 mL of DMSO were added into each vial containing the frozen filter. Vials were then incubated for 20 min at 65°C, then cooled in a dark box to room temperature (approximately 1 hr). Four mL of buffered Acetone (90% Acetone, 10% saturated MgCO_3_) were added to the vial and thoroughly mixed. Vials were then left to settle for few minutes and 3 mL sample was drown form the vial to a fluorometer cuvette. Fluorescence was measured with a calibrated Trilogy fluorometer (Turner Designs) using the non-acidification method (Welschmeyer and Naughton 1994).

### DNA extraction

The relative abundance of prominent microbial taxa (phylotypes) in the seawater was estimated using next-generation sequencing (NGS) of SSU rRNA genes to evaluate any differential removal by suspension feeders benthic community. Ten mL of seawater collected from each station and filtered on a 25 mm, 0.2 μm polycarbonate membrane (GE Healthcare Biosciences, cat. No. 110606) under low vacuum and frozen in 1.5 mL micro-tubes at -20°C until analysis. DNA from each filter was extracted using the DNeasy ‘blood & tissue kit’ (QIAGEN, Cat. No. 69504) with the following modifications to the manufacturer’s protocol: ATL buffer (180 μL) and 20 μL of proteinase K were added and samples were incubated at 56°C for 1 hr. Then 200 μL of AL Buffer and 200 μL of 95-100% ethanol was added to the sample and the mixture was pipetted into spin columns and placed in a 2 mL collection tube. Tubes were centrifuged at 6000 RCF for 1 min. The flow-through was discarded and 500 μL of AW1 buffer was added to the column, centrifuged at 6000 RCF for 1 min, and the flow-through again discarded. This step was repeated for the third time, with 500 μL Buffer AW2 and a spin of 18,000 RCF for 1 min to dry the membrane before elution. For the elution step, the spin column was placed on a new collection tube. Two hundred µL of buffer AE preheated to 56°C was pipetted at three steps (50 µL, 50 µL, and 100 µL) into the column and each step was followed by 6000 RCF centrifugation for 1 min. The sample was then incubated at room temperature for at least a minute and stored at -20°C.

### Next-generation sequencing

Samples were sequenced by Research and Testing Laboratories (Lubbock TX). The SSU rRNA genes were amplified for sequencing using a forward and reverse fusion primers (515F-Y - 926R (Parada et al. 2016). The forward primer was constructed with (5’-3’) the Illumina i5 adapter (AATGATACGGCGACCACCGAGATCTACAC), an 8-10bp barcode, a primer pad, and the 5’– GAGTTTGATCNTGGCTCAG –3’ primer. The reverse fusion primer was constructed with (5’-3’) the Illumina i7 adapter (CAAGCAGAAGACGGCATACGAGAT), an 8-10bp barcode, a primer pad, and the 5’– GTNTTACNGCGGCKGCTG –3’ primer. Primer pads were designed to ensure the primer pad/primer combination had a melting temperature of 63°C-66°C, according to methods developed by Patrick Schloss’ laboratory (http://www.mothur.org/w/images/0/0c/Wet-lab_MiSeq_SOP.pdf). Amplifications were performed in 25 μL reactions with Qiagen HotStar Taq master mix (Qiagen Inc, Valencia, California), 1μL of each 5 μM primers, and 1 μL of the template. Reactions were performed on ABI Veriti thermocyclers (Applied Biosystems, Carlsbad, California) under the following cycle conditions: 95°C for 5 min, then 35 cycles of 94°C for 30 sec, 54°C for 40 sec, 72°C for 1 min, followed by one cycle of 72°C for 10 min and a final 4°C hold.

Amplification products were visualized with eGels (Life Technologies, Grand Island, New York). Products were then pooled equimolarly and each pool was size-selected in two rounds using Agencourt AMPure XP (BeckmanCoulter, Indianapolis, Indiana) in a 0.7 ratio for both rounds. Size-selected pools were then quantified using a Qubit 2.0 fluorometer (Life Technologies) and loaded on an Illumina MiSeq (Illumina, Inc. San Diego, California) 2×300 flow cell at 10 pM.

### Sequence data analysis

SSU rRNA sequences were treated using a qiime2 (v. 2018-8) and biom-format (v. 2.1.6) deployed through a bash scripts (Suppl. File). Briefly demultiplexed forward and reverse reads were imported into qiime2 artifacts and ASV tables were generated using *qiime dada2* using options “--p-trim-left-f 19 --p-trim-left-r 20 --p-trunc-len-f 300 --p-trunc-len-r 250” The resulting ASVs were identified using *qiime feature-classifier* using an in house version of the Silva132 (arb-silva.de) including only the region flanked by the primers, and a taxonomy files extracted from this database (see data availability below for details).

Since the primers we used amplify both 16S rRNA from prokaryotes and chloroplasts as well as 18S rRNA from eukaryotes, the ASV table and sequence files were filtered using *qiime taxa* to generate three sets of table/sequences, one with 16S rRNA, one with chloroplasts and one with eukaryotic sequences. ASV tables were modified to include taxonomy using *sed* and *gawk* and *biom-format* and exported in “.tsv” format for further analyses. Here we describe only results concerning 2388 prokaryotic ASVs.

### Data analysis

Due to temporal changes in microbial communities, and assuming the entire transect represented a single water mass the concentrations along each transect were normalized to the seaward-most station and presented as “% of open sea”. The significance level of cross-shore trends was tested using the “Page test” for ordered alternatives (Page 1963). This non-parametric test is a modified version of the Kruskal-Wallis one-way ANOVA for ranked data. Nearshore depletion of microbial taxa was tested (each taxon per season) with H_1_ as an ordered decrease in concentrations from the “open water” toward the shore. Due to missing sampling points, only 3 complete transects toward the hard-bottom were used for the test in each season.

Data collected with the CTD was converted, aligned, and binned (at 0.1 m) with the SBE DataProcessing software (Version 7.2). Since salinity differences along the transects were negligible (<0.01 PSU), density differences were driven solely by temperature. The vertical and horizontal change in temperature and *in-vivo* chlorophyll fluorescence along the cross-shore transects (hereafter ‘anomaly’) was calculated within each transect as anomalies (the difference of each data point from the average temperature or fluorescence along the transect). All seasonal anomalies were plotted together using Ocean Data View (Version 5). Interpolation was made with the Weighted-average gridding function of Ocean Data View using a seeking distance of 0.25 m along the vertical axis and 100 m along the horizontal axis. Due to the low (N=4), data are presented as a seasonal average ± standard error (SE) unless otherwise indicated.

Removal of specific ASVs toward the shore was calculated by multiplying the relative abundance of each ASV and the total bacterial cell counts obtained by flow cytometry and then calculation as a percentage of the open sea station. Normalized removal was calculated by normalizing measured removal to the ASV with the highest removal within the same transect. Implementation of this approach provides a powerful tool to indirectly evaluate the effect of benthic removal of microbial prey at the ASV level. Hereafter, the use of the terms “selectivity” and “preference” are limited to their technical definition (Chesson 1978, 1983), i.e., the removal of a prey type in higher proportion than its proportional presence in the environment, relative to other food types present.

### Data availability

Raw sequencing reads are available through the NCBI SRA under accession number PRJNA912166. The analysis pipeline and associated files including scripts, mapping files, taxonomic identification databases and ASV tables are available through a github repository (github.com/suzumar/transect_ms).

## Results

During winter, a temperature gradient was found along the transects with colder water at the shallow stations above the hard-bottom (stations 1-5, 19.4±0.03°C, Fig 2a) in comparison to the “open sea” (stations 6-8, 20.1±0.03°C, Fig 2a). During spring, no such temperature gradient was observed along the transects but the beginning of stratification was noticeable, with a half-degree Celsius warmer surface layer (0-3 m, 20.1°C±0.07, Fig 2b) than the deeper water (3-12m, 19.6°C±0.02, Fig 2b). The warmer, nearshore surface layer (10-25 m depth, Fig S1) showed typical coastal enrichment, with higher chlorophyll concentrations (Fig S1) and cell counts (Fig S2) than the open sea. Salinity was similar along the transect (1.5m depth) and with depth (0-12m) during winter and spring (39.26-39.31 PSU and 39.35-39.28 PSU, respectively, during winter and, 39.29-39.23 PSU and 39.06-39.21 PSU, respectively, during spring).

**Figure 2:**
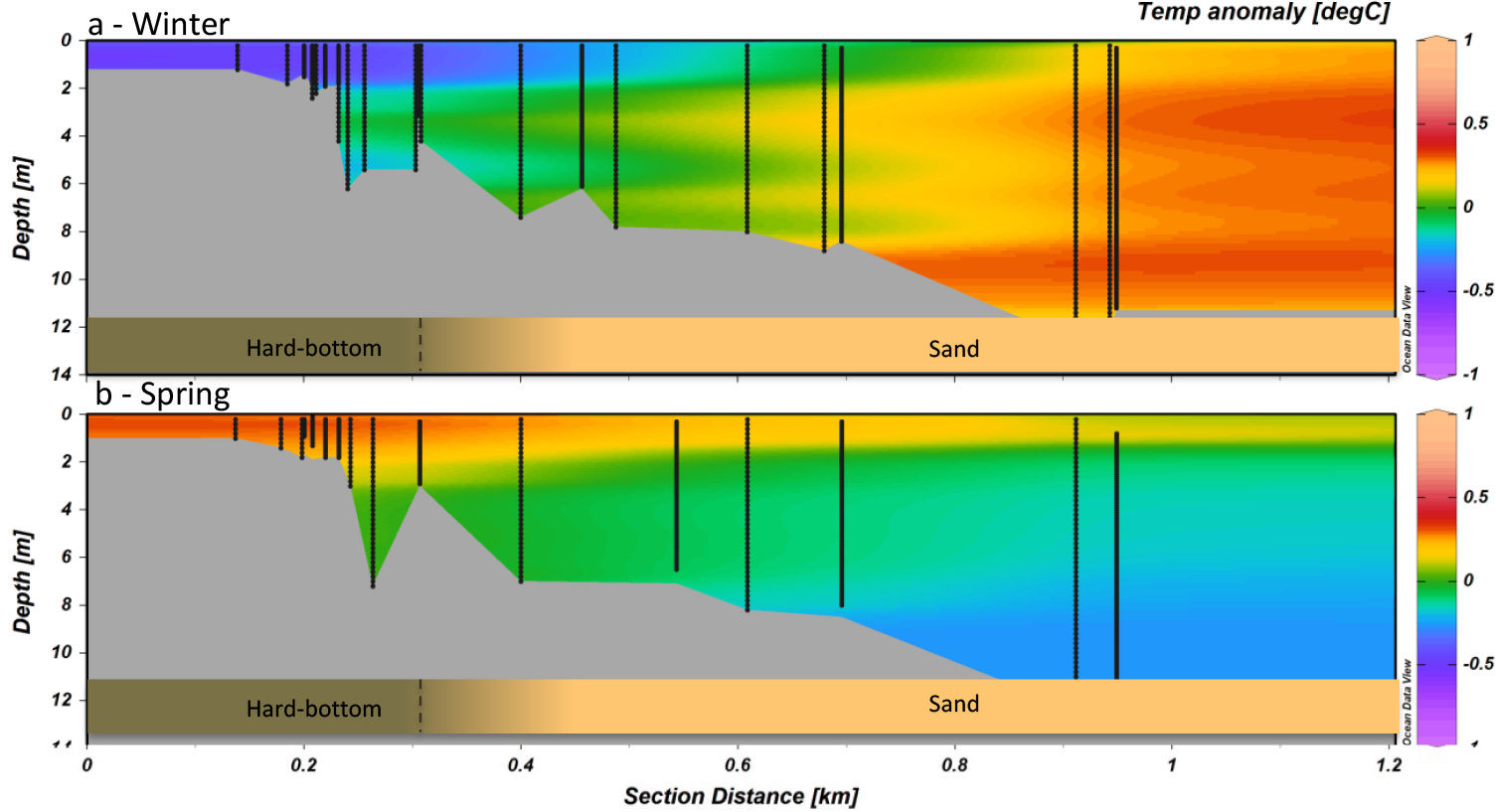
Temperature anomaly along the cross-shore transects. (a) Winters of 2015-2016, n=4. (b) Springs of 2016-2017, n=4. Sampling stations are shown as vertical black lines. Grey shading approximates the bottom depths based on the bottom depth of each cast. In most cases, the CTD was lower all the way to the bottom (∼0.2 mab).

In the cross-shore transects, the concentration of all microbial populations decreased toward shore in a highly significant gradient (Page test, p<0.01, Fig. 3a-d). In most cases, and for most of the microbial populations, cell concentrations were similar along the ∼700 m sandy section of the transects, between the open sea and the outer boundary of the rocky area (Stations 6 to 8). Sometimes even an increase was observed along that section (Table S1, S2). Most of the depletion occurred above the rocky habitat (Stations 1 to 5). The depletion of the microbial populations over the rocky section range between 25-50% of their percentage of the open sea (Fig 3a-d). *In vivo* chlorophyll fluorescence (Fig S3) and the concentration of extracted chlorophyll a also showed a strong depletion above the hard-bottom area compared to the open sea (Fig 3e).

**Figure 3:**
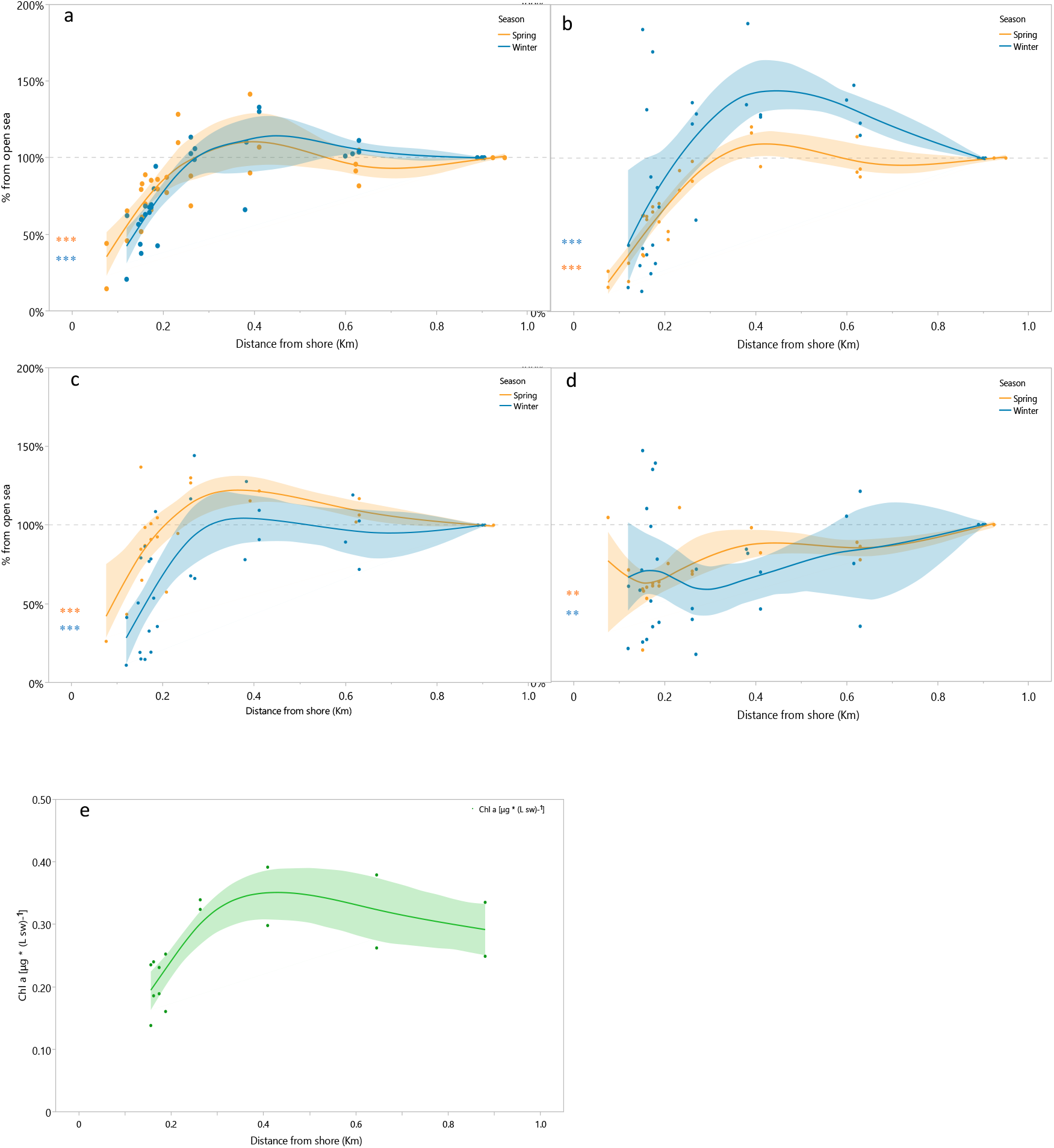
Cross-shore transects showing the percent change in the concentrations of several microbial cell populations (counted by flow cytometry) from their “open sea” concentration ∼900 m offshore. (a) Bact, non-photosynthetic bacteria, (b) Euk, eukaryotic algae, (c) Syn, *Synechococcus sp*. (d) PLP, *Prochlorococcus-* like particles, (e) Extracted chlorophyll *a* concentration along 3 transects. Blue, n=4, transects performed during winter. Orange, n=4, transects performed during spring. The significance of the Page test for a gradient of decreasing concentrations toward the shore is denoted as **<0.01, ***<0.001

Differential changes in relative abundances between ASVs and even within the same ASV in different transects were observed (Fig 4a-c). For example, ASVs belonging to cyanobacteria group were on average 8%, 35% and 3% of their percentage at the open sea (Fig 4a-c accordingly).

**Figure 4:**
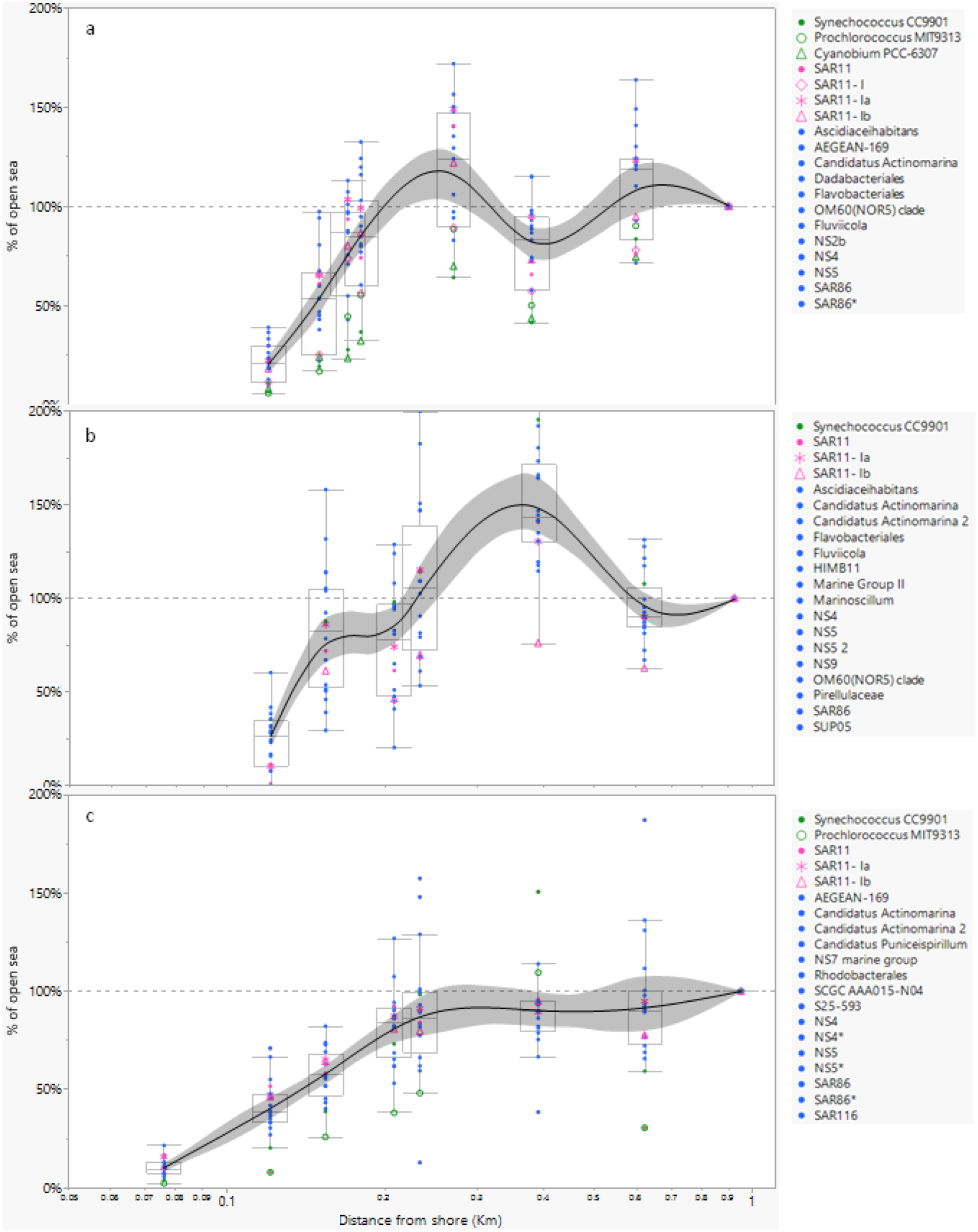
Cross-shore transects showing the percent change in the concentrations of the 20 most abundant prokaryotic ASV s in the water relative to their ‘open sea’ concentration ∼900 m offshore: (a-c) transects above rocky hard bottom (January 2016, March 2016 and, April 2016). The vertical dashed line represents the concentration in the open sea (100%). Pink indicates members of SARI 1 clade, green indicates autotrophs, and blue indicates other non-photo synthetic bacteria.

Within the SAR11 clade, the percentage of four ASVs decreased along the transects, in similar trends (Fig 5). Transects showed 83% ±7% (Mean ±SD, Fig 6a-c) normalized removal for the 20 most abundant ASVs. Abundant ASVs including *Synechococcus* CC9901 (27% relative abundance, transect during January 2016) and SAR11-Ia (25%, transect during April 2016) were removed at similar percentages (95% and 84% accordingly) compared to less abundant ASVs (e.g. SAR11-Ib 1.3% relative abundance and 87% removal, SAR86 2.3% relative abundance and 86% removal) (Fig 6).

**Figure 5:**
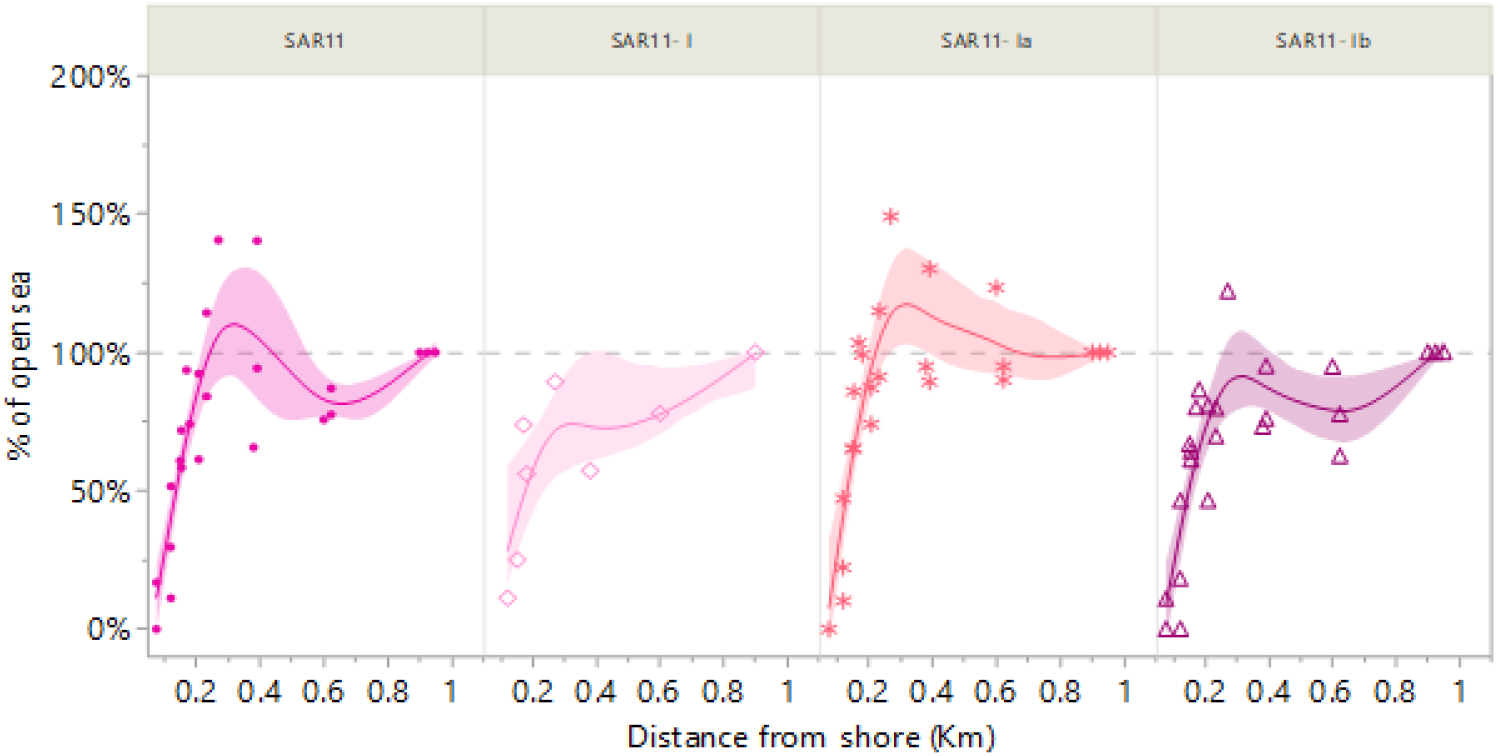
Cross-shore transects showing the percent change in the concentrations of four ASVs belonging to SAR11 clade in the water relative to their “open sea” concentration ∼900 m offshore, (a-c) transects above rocky hard bottom (January 2016, March 2016 and, April 2016). The vertical dashed line represents the concentration in the open sea (100%).

**Figure 6:**
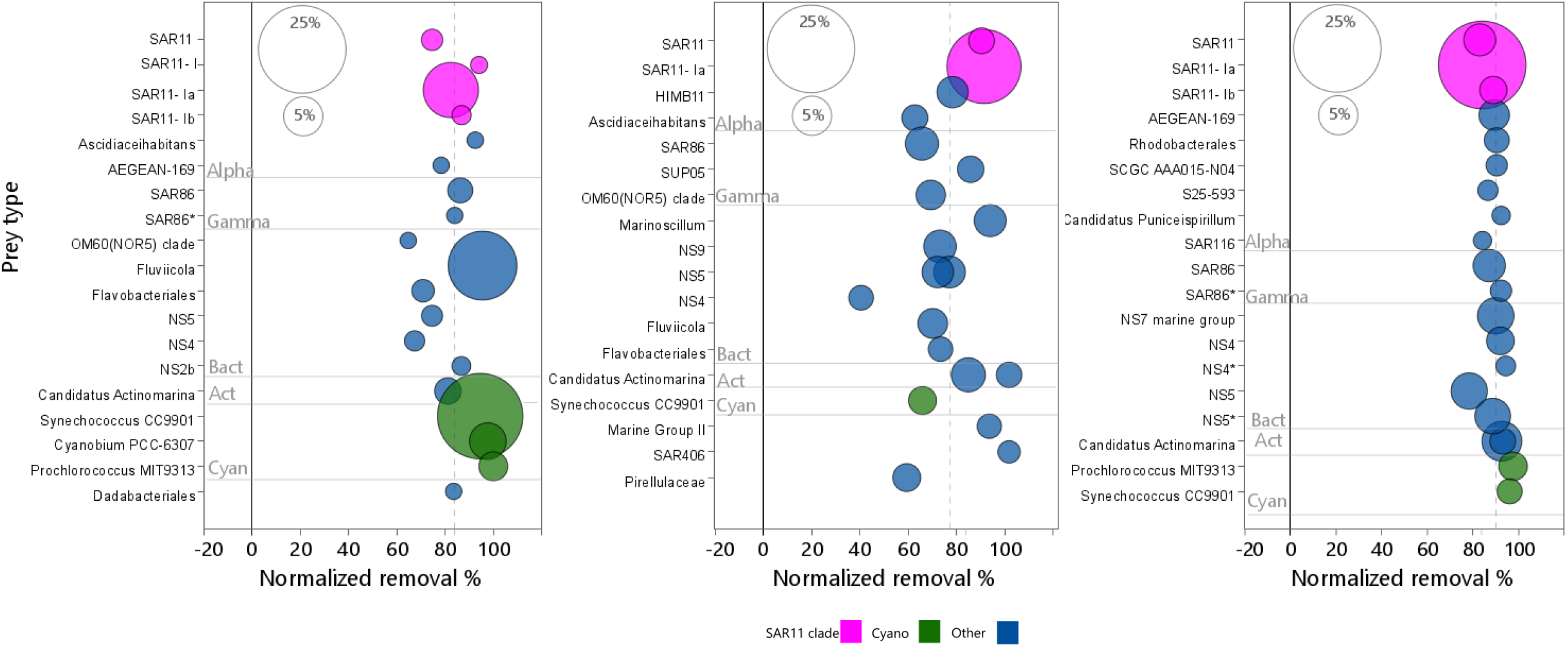
Normalized removal of the 20 most abundant ASVs from the open sea toward the shore. (a-c) transects above rocky hard bottom (January 2016, March 2016 and, April 2016). Pink indicates members of SAR11 clade, green indicates autotrophs, and blue indicates other non-photosynthetic bacteria. Dashed vertical line represents the expected retention assuming equal retention probability for all cells. Size of circles represents relative abundance in the open water (Station 8) during sampling., with the two white circles plotted for scaling.

## Discussion

This study was designed to evaluate whether the effect of differential removal of microorganisms by benthic suspension feeders can be detected at the community level. Most of the microbial populations were depleted above the hard-bottom area compared to the open sea. Differential removal was observed at low extent (between and within species). A similar trend was also observed for chlorophyll *a* concentration, suggesting the formation of a depleted boundary layer over the hard-bottom. The hard-bottom community at the study site included a diverse assemblage of suspension feeders, including sponges, bivalves, ascidians, polychaetes, hydrozoans, and bryozoans (Rilov et al. 2018). Suspensions feeders on the hard-bottom occur in different densities, and hence the competition for available prey is between species, within species and even between different taxa. Niche speciation is expected in such diverse and dense community where different taxa utilize different filtration mechanisms and presumably, different organic carbon sources in their diet. If the depletion of microbial cells is the outcome of grazing by benthic suspension feeders, near-bottom depletion should generate a shore-wise depletion above the hard-bottom section of the transects compared to the sandy section of the transect (e.g., Genin et al. 2009; Jones et al. 2009). In case other processes determine the cross-shore trend (e.g., runoff, eutrophication, and coastal pollution), the depleted zone is expected to extend throughout the water column and no difference is expected along the transects. Differential grazing of phytoplankton by dense populations of benthic suspension feeders was also reported in San-Francisco bay where it was attributed to different sinking rates of the microalgae (Lucas et al. 2016). Such a phenomenon is very unlikely in the east Mediterranean where the planktonic community is dominated by very small cells (<10 μm) with negligible sinking rates (Siokou-Frangou et al. 2010). This body of prior work supports the assumption that the depletion reported here is due to benthic suspension grazing although it was not measured directly.

Physical and biological processes are the major factors controlling changes in particle concentrations throughout the ocean, and advective processes could be responsible for the pico and nanoplanktom depletion we observed. Monismith et al. (2006) showed that shallow regions nearshore experience larger temperature changes than deeper regions offshore. When the water warms during the daytime (e.g., in the spring), the shallow near-shore water body tends to warm faster than the nearby open sea. The warmer water expands and flows offshore, causing deeper and cooler water to flow onshore at depth to replace it. This ‘thermal flow’ (Fig 7) leads to an upwelling of deeper water and material to the nearshore region (Fig 7). The opposite process occurs during winter or cold nights, when a faster cooling of the shallow, near-shore waterbody initiates near-shore downwelling of cold surface water that reverses the direction of the “thermal flow” cycle and induces onshore flow of surface water (Fig 7). In the Eastern Mediterranean Sea, the water column is usually stably stratified, and the numbers and biomass of surface water plankton is lower than subsurface layers (Suari et al 2019). Onshore transport of surface water from the offshore during the cold phase of the thermal flow cycle and the nearshore downwelling of “plankton poor” seawater is expected to yield a benthic zone of depleted of plankton close to shore (Labiosa et al. 2003). However, this physical mechanism would not be expected to change the relative abundance of different microbial groups along the transects, nor should it differentially affect rocky and sandy sections of the transects. Moreover, during the warm phase of the thermal flow cycle (in spring for example), local upwelling is expected to bring plankton-rich water onshore, countering removal, and the benthic zone should be as rich if not richer nearshore. Our results showed that nearshore depletion occurred during both the cold and hot ‘phase’ of the thermal flow, suggesting that a biological process such as filtration is probably the mechanism responsible for the depletion above the hard-bottom section in the transects rather than advection of planktonic poor water. A similar conclusion was reached by Patten et al. (2011), who showed depleted levels (∼40% on average) of microbial cells over a reef with negligible removal over a sandy bottom.

**Figure 7:**
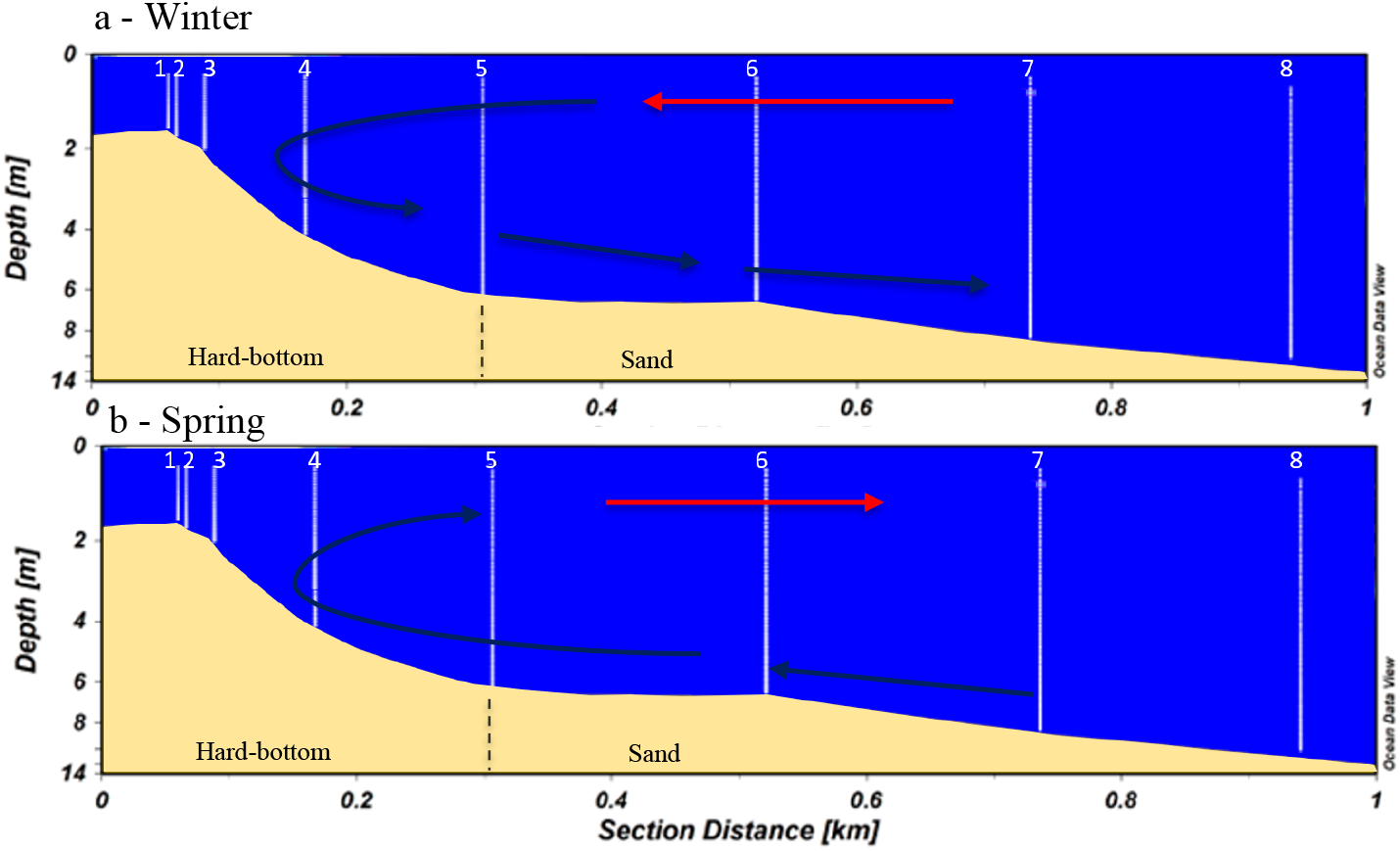
Schematic illustration of a near-shore thermal flow cycle. (a) during winter, when cooling of nearshore water drives offshore flow at depth and onshore flow at the surface. (b) during spring, when the shallow water near shore warms and water flow offshore at the surface and onshore below.

Our previous work showed partial grazing resistance of members SAR11 clade to grazing by pelagic and benthic tunicates (Dadon-Pilosof et al. 2017), and in general, lower retention efficiencies on LNA bacteria than other prey types. Selective grazing and specifically low retention on LNA compare to both HNA bacteria and *Synechococcus* was also observed in sponges (Hanson et al. 2009; Jiménez 2011) and preferential retention of *Synechococcus* and eukaryotic algae over other prey types was observed in bivalves (Yahel et al. 2009). The results (although using indirect measurements) of this study suggested inversely to what we expected that members of SAR11 were grazed at higher efficiencies than other available prey (Fig 5,6) somewhat reflecting their overall high abundance. This work strengthens the niche speciation assumption, hinting that some taxa within the hard-bottom community specialize on grazing SAR11. Besides the obvious members of the suspension feeding community, we must consider the heterotrophic nanoflagellates attached to the rocks (Yahel et al. 2006). This cryptic community has a large grazing effect that may be part of the explanation for prey depletion above the hard-bottom. It is known that heterotrophic nanoflagellates are important bacterivores in pelagic waters (Tophøj et al. 2018).

The apparent lack of grazing evasion by members of SAR11 could be possibly explained by a different mode of interaction to other filter feeders compared to tunicates. Suspension feeders have different filtration organs, using cilia, mucus, or both to capture and process suspended particles. While some are active suspension feeders and specialize in filtration of small particles, others are passive feeders and specialize in filtration of large particles (Gili and Coma 1998). Different grazing strategies of benthic suspension feeders would potentially increase the opportunities for exploitation of available prey by communities of suspension feeders (Gili and Coma 1998). Specialization of different suspension feeders on different prey would explain the homogeneous decrease of different available prey. While removal by benthic filter feeders is a likely biological explanation for the depletion of pico- and nanoplankton we observed, other processes such adsorption to mucus might also have contributed (Decho 1990) although members of SAR11 clade are known as free-living and not particle-associated or biofilm forming bacteria (Giovannoni 2017, Haro-Moreno et al. 2020)

Evaluation of evasiveness using “in versus out” experiments such those in Dadon-Pilosof et al. (2017) with other taxa, or the measurement of adsorption of different bacteria including cultured *Candidatus* Pelagibacter ubique to transparent exopolymer particles (Long and Azam 1996) with water from different points in the transect remain as possible future experiments to further clarify our results.

Measurements of removal rates of microbial plankton by benthic organisms at the level of the whole community remain challenging despite of decades of studies of this theme (Sargent and Austin 1949; Odum and Odum 1955; Johannes et al. 1972). More recent studies, assessed spatial gradients of DOC, bacterioplankton and virioplankton concentrations in reef ecosystem resulting in depleted microbial community over the reef compare to negligible removal over sandy bottom nearby (Patten et al. 2011; Nelson et al. 2011). An alternative approach utilizes a control-volume approach either by physically enclosing the community in a bell jar for in situ measurements (e.g., (Hopkinson et al. 1991) or by enclosing and imaginary “box” over the bottom using vertical arrays of samplers and ADCPs to quantify the fluxes of plankton into and out of the control volume (Genin et al. 2002, 2009). A third approach integrates individual rate measurements of dominant benthic suspension feeders with their abundance and size distribution to assess the community flux and its effect on the planktonic community in the overlying water (e.g., Genin et al. 2009; Lucas et al. 2016 and references therein).

Further investigation is required to develop and estimate, based on indirect evaluation, cells removal above hard-bottom suspension feeders. Methodology fine tuning is required for estimating the cells removal above the hard-bottom compared to open sea and to discriminate grazing, other biological processes and physical mechanisms. Previous studies showed that SAR11 evade predation of benthic and pelagic ascidians mucus, but this study showed (albeit indirectly) that SAR11 was removed efficiently as other prey cells by hard bottom grazers.

Potential grazers might be sponges, bivalves, nanoflagellates attached to the surface of the rocks which filter their prey by using several different mechanisms. Understanding the mechanisms and the variance between the grazers’ preferences and/or the microbial cells abilities to evade predation has ramifications on processes affecting the marine food web such as top-down control and, benthic pelagic coupling. Grazing resistance mechanisms are still understudied and should be investigated further to gain knowledge on its effects in the marine ecosystem.

## Supporting information

Supplementary material_Significant SAR11 removal

## Acknowledgements

Research funding was provided by Ruppin Academic center, support provided to A.D.P by the Mediterranean Sea Research Center of Israel. I thank my supervisors Prof. Amatzia Genin and Prof. Gitai Yahel for their advice along the way. Special gratitude to R. Rosenblatt for availability and technical assistance throughout the field experiments.

## Author Contributions

**A.D.P**. designed the study and participated in field experiments, data analysis, and manuscript preparation. **K.R.C**, participated in data analysis, and contributed substantially to drafting the manuscript. **M. T.S** designed and performed bioinformatics analyses and contributed substantially to drafting the manuscript. All authors discussed the results and commented on the manuscript during its preparation and approved the submitted version of the manuscript.

## Notes

### Competing Interest Statement

The authors have declared no competing interest.

