## Supplementary material_Significant SAR11 removal for "Significant SAR11 removal by a hard-bottom community"

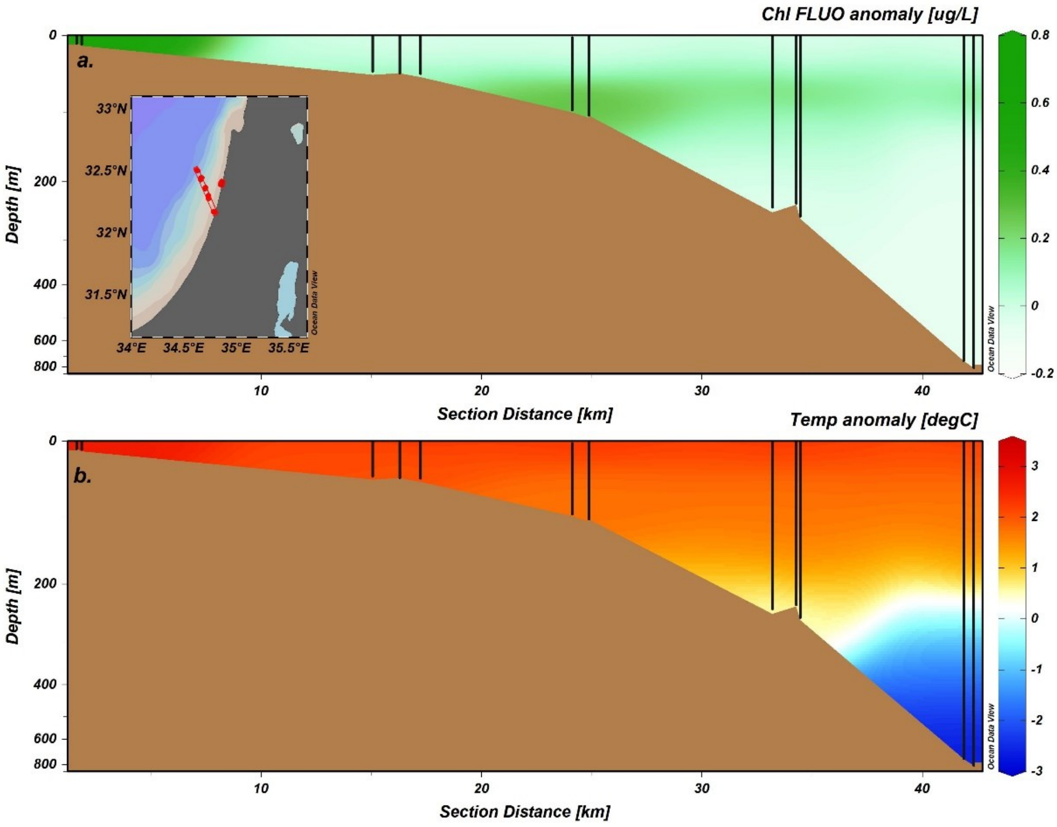


Figure S1: Cross-shelf transect across the Israeli shelf at the eastern Mediterranean Sea measured during March 2018. Sampling stations are shown as vertical black lines. Brown shading approximates the bottom based on the logged bottom depths at the sampling locations. (a) chlorophyll fluorescence (b) Temperature. Interpolation is based on the weighted-average gridding algorithm of Ocean Data View (Ver. 5.12) with a horizontal seeking distance of 3 km and a vertical seeking distance of 72 m. Note that the upper part of y-axis is stretch so that changes in the surface layer are more pronounced.

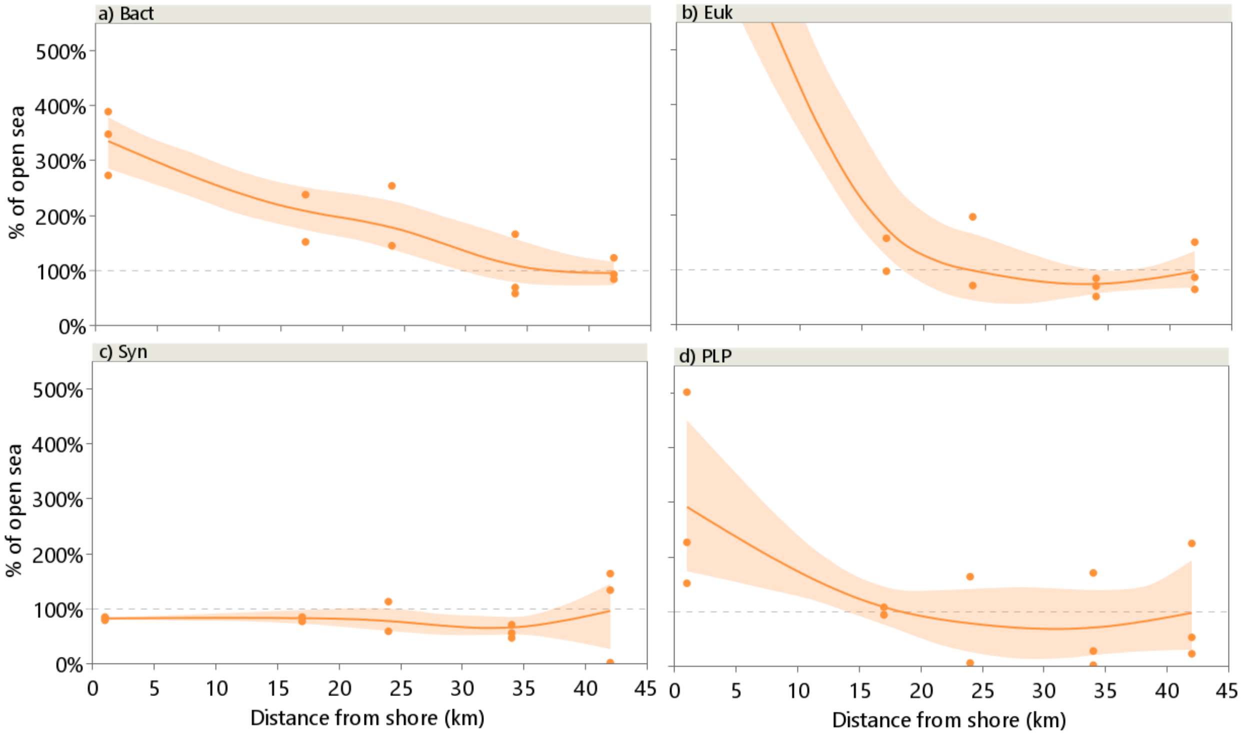


Figure S2: Cross-shore transects showing the percent change in the concentrations of several microbial cell populations (counted by flow cytometry) from their open sea concentration 42 km offshore as sampled in the Eastern Mediterranean Sea, March 2018 (n=3). (a) Bact, non-photosynthetic bacteria, (b) Euk, eukaryotic algae, (c) Syn, *Synechococcus sp.* (d) PLP, *Prochlorococcus*- like particles.

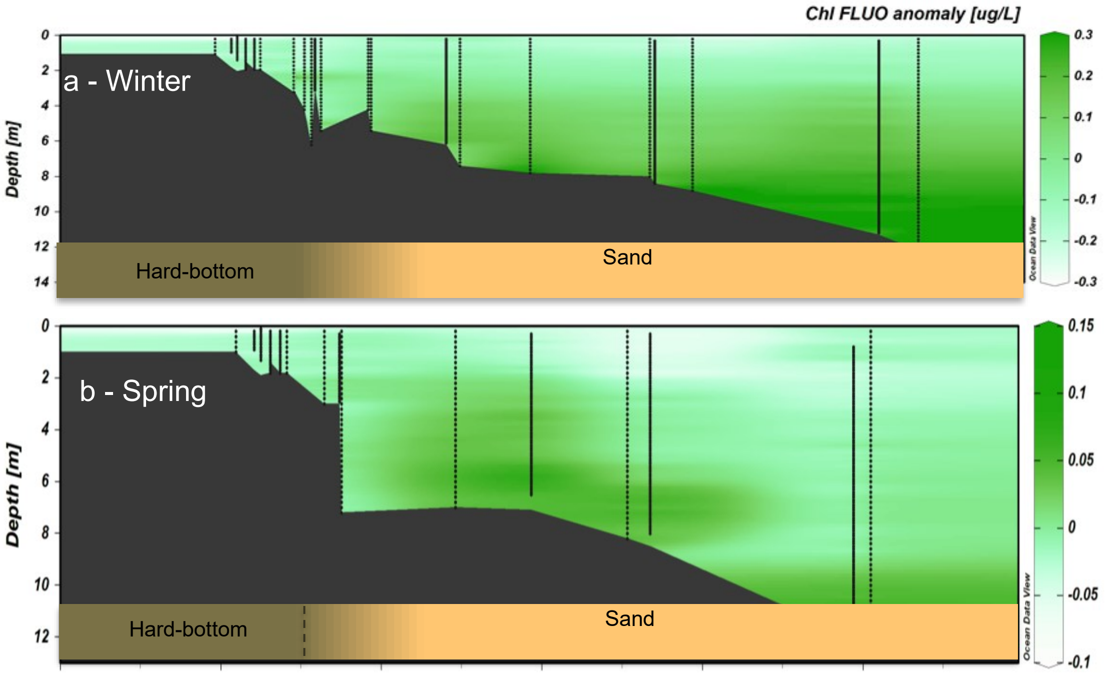


Figure S3: In-vivo chlorophyll fluorescence anomaly along the cross-shore transects. (a) Winters of 2015-2016, n=4. (b) Springs of 2016-2017, n=4. All other details as in Fig. 2.

Table S1: Summary of all cell-counts for all transects. Euk, eukaryotic algae, Syn, *Synechococcus sp.,* PLP, *Prochlorococcus*- like particles*,* Bact, non-photosynthetic bacteria

| **Transect details** | **Station No.** | **Distance from shore (m)** | **Euk cells mL$^{-1}$** | **Syn cells mL$^{-1}$** | **PLP cells mL$^{-1}$** | **Bact cells mL$^{-1}$** |
|---|---|---|---|---|---|---|
| 27/12/2015 Hard-bottom | 1 | 121 | 6.74x $10^2$ | 1.43x $10^4$ | 1.77x $10^4$ | 3.94x $10^5$ |
| | 2 | 146 | 4.67x $10^2$ | 2.02x $10^4$ | 1.68x $10^4$ | 3.35x $10^5$ |
| | 3 | 170 | 1.37x $10^3$ | 2.88x $10^4$ | 2.08x $10^4$ | 4.73x $10^5$ |
| | 4 | 184 | 1.26x $10^3$ | 4.09x $10^4$ | 3.05x $10^4$ | 6.06x $10^5$ |
| | 5 | 269 | 9.31x $10^2$ | 4.49x $10^4$ | 3.40x $10^4$ | 6.78x $10^5$ |
| | 6 | 383 | 2.93x $10^3$ | 4.61x $10^4$ | 3.49x $10^4$ | 6.73x $10^5$ |
| | 7 | 616 | 2.31x $10^3$ | 4.39x $10^4$ | 3.17x $10^4$ | 6.87x $10^5$ |
| | 8 | 905 | 1.56x $10^3$ | 4.09x $10^4$ | 3.01x $10^4$ | 6.60x $10^5$ |
| 17/01/2016 Hard-bottom | 1 | 120 | 1.13x $10^3$ | 7.45x $10^3$ | 2.88x $10^3$ | 1.88x $10^5$ |
| | 2 | 150 | 9.51x $10^2$ | 1.40x $10^4$ | 7.82x $10^3$ | 3.63x $10^5$ |
| | 3 | 170 | 1.79x $10^3$ | 2.31x $10^4$ | 1.36x $10^4$ | 5.41x $10^5$ |
| | 4 | 180 | 2.27x $10^3$ | 3.98x $10^4$ | 1.83x $10^4$ | 6.85x $10^5$ |
| | 5 | 270 | 9.36x $10^3$ | 4.61x $10^4$ | 1.57x $10^4$ | 9.12x $10^5$ |
| | 6 | 380 | 9.79x $10^3$ | 6.14x $10^4$ | 1.16x $10^4$ | 5.92x $10^5$ |
| | 7 | 600 | 1.00x $10^4$ | 6.99x $10^4$ | 1.45x $10^4$ | 8.96x $10^5$ |
| | 8 | 900 | 7.27x $10^3$ | 7.88x $10^4$ | 1.37x $10^4$ | 9.01x $10^5$ |
| 24/03/2016 Hard-bottom | 1 | 76 | 7.50x $10^2$ | 4.48x $10^3$ | 3.15x $10^3$ | 3.06x $10^5$ |
| | 2 | 121 | 9.00x $10^2$ | 7.84x $10^3$ | 5.42x $10^3$ | 4.64x $10^5$ |
| | 3 | 154 | 1.79x $10^3$ | 1.16x $10^4$ | 7.84x $10^3$ | 5.90x $10^5$ |
| | 4 | 208 | 1.49x $10^3$ | 1.06x $10^4$ | 7.22x $10^3$ | 5.32x $10^5$ |
| | 5 | 233 | 2.63x $10^3$ | 1.73x $10^4$ | 1.20x $10^4$ | 7.79x $10^5$ |
| | 6 | 391 | 3.45x $10^3$ | 2.11x $10^4$ | 1.37x $10^4$ | 9.45x $10^5$ |
| | 7 | 623 | 2.60x $10^3$ | 1.82x $10^4$ | 1.11x $10^4$ | 4.77x $10^5$ |
| | 8 | 924 | 2.86x $10^3$ | 1.80x $10^4$ | 1.09x $10^4$ | 6.97x $10^5$ |
| 24/03/2016 Sand | 1 | 61 | 4.96x $10^3$ | 1.39x $10^4$ | 5.15x $10^4$ | 5.46x $10^5$ |
| | 2 | 118 | 3.76x $10^3$ | 1.61x $10^4$ | 2.18x $10^4$ | 5.06x $10^5$ |
| | 3 | 153 | 3.19x $10^3$ | 1.70x $10^4$ | 1.72x $10^4$ | 5.01x $10^5$ |
| | 4 | 248 | 2.67x $10^3$ | 2.25x $10^4$ | 1.15x $10^4$ | 4.44x $10^5$ |
| | 5 | 326 | 2.59x $10^3$ | 1.79x $10^4$ | 1.09x $10^4$ | 5.54x $10^5$ |
| | 6 | 560 | 3.41x $10^3$ | 1.98x $10^4$ | 1.18x $10^4$ | 4.38x $10^5$ |
| | 7 | 623 | NA | NA | NA | NA |
| | 8 | 924 | NA | NA | NA | NA |
| 19/04/2016 Hard-bottom | 1 | 76 | 2.09x $10^2$ | 1.79x $10^3$ | 3.72x $10^2$ | 7.29x $10^4$ |
| | 2 | 121 | 2.59x $10^2$ | 1.38x $10^3$ | 5.57x $10^2$ | 2.37x $10^5$ |
| | 3 | 154 | 4.82x $10^2$ | 3.27x $10^3$ | 4.83x $10^2$ | 3.18x $10^5$ |
| | 4 | 208 | 6.22x $10^2$ | 4.05x $10^3$ | 6.36x $10^2$ | 4.52x $10^5$ |
| | 5 | 233 | 1.05x $10^3$ | 5.96x $10^3$ | 1.12x $10^3$ | 6.43x $10^5$ |
| | 6 | 391 | 1.54x $10^3$ | 1.03x $10^4$ | 5.67x $10^2$ | 4.68x $10^5$ |
| | 7 | 623 | 1.51x $10^3$ | 9.70x $10^3$ | 5.64x $10^2$ | 4.94x $10^5$ |
| | 8 | 950 | 1.33x $10^3$ | 5.78x $10^3$ | 5.05x $10^2$ | 5.16x $10^5$ |

| Transect details | Station No. | Distance from shore (m) | Euk cells $mL^{-1}$ | Syn cells $mL^{-1}$ | PLP cells $mL^{-1}$ | Bact cells $mL^{-1}$ |
|---|---|---|---|---|---|---|
| 05/12/2016 Hard-bottom | 1 | 152 | $2.88x 10^2$ | $5.22x 10^3$ | $6.94x 10^2$ | $1.76x 10^5$ |
| | 2 | 161 | $2.59x 10^2$ | $5.46x 10^3$ | $6.39x 10^2$ | $2.97x 10^5$ |
| | 3 | 174 | $3.03x 10^2$ | $7.37x 10^3$ | $8.17x 10^2$ | $3.17x 10^5$ |
| | 4 | 188 | $4.77x 10^2$ | $9.89x 10^3$ | $4.05x 10^2$ | $2.00x 10^5$ |
| | 5 | 261 | $9.55x 10^2$ | $1.72x 10^4$ | $6.82x 10^2$ | $5.34x 10^5$ |
| | 6 | 411 | $8.99x 10^2$ | $2.36x 10^4$ | $1.28x 10^3$ | $6.15x 10^5$ |
| | 7 | 630 | $8.05x 10^2$ | $2.93x 10^4$ | $9.07x 10^2$ | $5.27x 10^5$ |
| | 8 | 890 | $7.01x 10^2$ | $2.71x 10^4$ | $1.14x 10^3$ | $4.70x 10^5$ |
| 08/12/2016 Hard-bottom | 1 | 152 | $2.20x 10^3$ | $1.78x 10^4$ | $1.19x 10^4$ | $2.58x 10^5$ |
| | 2 | 161 | $1.58x 10^3$ | $1.80x 10^4$ | $8.17x 10^3$ | $2.96x 10^5$ |
| | 3 | 174 | $2.03x 10^3$ | $1.77x 10^4$ | $9.20x 10^3$ | $3.00x 10^5$ |
| | 4 | NA | NA | NA | NA | NA |
| | 5 | 261 | $1.47x 10^3$ | $2.63x 10^4$ | $3.11x 10^3$ | $4.48x 10^5$ |
| | 6 | 411 | $1.52x 10^3$ | $2.49x 10^4$ | $3.16x 10^3$ | $5.80x 10^5$ |
| | 7 | 630 | $1.47x 10^3$ | $1.50x 10^4$ | $3.50x 10^3$ | $4.53x 10^5$ |
| | 8 | 890 | $1.20x 10^3$ | $2.22x 10^4$ | $7.49x 10^3$ | $4.36x 10^5$ |
| 19/04/2017 Hard-bottom | 1 | 152 | $6.90x 10^2$ | $5.44x 10^3$ | $2.46x 10^3$ | $2.34x 10^5$ |
| | 2 | 161 | $1.12x 10^3$ | $4.89x 10^3$ | $4.02x 10^3$ | $3.20x 10^5$ |
| | 3 | 174 | $1.21x 10^3$ | $4.95x 10^3$ | $4.93x 10^3$ | $3.00x 10^5$ |
| | 4 | 188 | $1.09x 10^3$ | $5.02x 10^3$ | $5.10x 10^3$ | $3.63x 10^5$ |
| | 5 | 261 | $1.58x 10^3$ | $8.25x 10^3$ | $3.86x 10^3$ | $3.12x 10^5$ |
| | 6 | NA | NA | NA | NA | NA |
| | 7 | 630 | $1.73x 10^3$ | $6.04x 10^3$ | $6.55x 10^3$ | $3.73x 10^5$ |
| | 8 | 890 | $1.87x 10^3$ | $4.59x 10^3$ | $8.32x 10^3$ | $4.52x 10^5$ |
| 19/04/2017 Hard-bottom | 1 | 152 | $1.14x 10^3$ | $5.42x 10^3$ | $5.22x 10^3$ | $3.82x 10^5$ |
| | 2 | 161 | $1.13x 10^3$ | $5.55x 10^3$ | $5.36x 10^3$ | $4.36x 10^5$ |
| | 3 | 174 | $1.25x 10^3$ | $5.85x 10^3$ | $5.55x 10^3$ | $4.16x 10^5$ |
| | 4 | 188 | $1.28x 10^3$ | $5.95x 10^3$ | $5.62x 10^3$ | $4.21x 10^5$ |
| | 5 | 261 | $1.79x 10^3$ | $1.35x 10^4$ | $9.70x 10^2$ | $4.26x 10^5$ |
| | 6 | 411 | $1.72x 10^3$ | $8.04x 10^3$ | $6.91x 10^3$ | $5.22x 10^5$ |
| | 7 | 630 | $1.60x 10^3$ | $6.78x 10^3$ | $6.65x 10^3$ | $5.12x 10^5$ |
| | 8 | 890 | $1.83x 10^3$ | $6.35x 10^3$ | $8.68x 10^3$ | $4.92x 10^5$ |
| 19/04/2017 Sand | 1 | 152 | $2.60x 10^3$ | $7.56x 10^3$ | $1.13x 10^4$ | $6.05x 10^5$ |
| | 2 | 161 | $2.48x 10^3$ | $6.96x 10^3$ | $1.14x 10^4$ | $6.46x 10^5$ |
| | 3 | 174 | $2.57x 10^3$ | $6.37x 10^3$ | $1.07x 10^4$ | $6.35x 10^5$ |
| | 4 | 188 | $2.04x 10^3$ | $7.18x 10^3$ | $1.01x 10^4$ | $4.81x 10^5$ |
| | 5 | 261 | $1.89x 10^3$ | $1.01x 10^4$ | $6.45x 10^3$ | $4.01x 10^5$ |
| | 6 | 411 | $2.00x 10^3$ | $6.78x 10^3$ | $7.77x 10^3$ | $4.98x 10^5$ |
| | 7 | 630 | $1.93x 10^3$ | $5.96x 10^3$ | $6.89x 10^3$ | $4.86x 10^5$ |
| | 8 | 890 | $1.61x 10^3$ | $3.68x 10^3$ | $6.43x 10^3$ | $4.62x 10^5$ |

Table S2: Average cell-counts by seasons. Results present as percentages of concentration measured for each group compared to the concentration measure at open sea, average± standard error of each station per season. Euk, eukaryotic algae, Syn, *Synechococcus sp.*, PLP, *Prochlorococcus*- like particles, Bact, non-photosynthetic bacteria

| Season (# of transects) | Station No. | Average Distance from shore (m) | Euk cells $mL^{-1}$ | Syn cells $mL^{-1}$ | PLP cells $mL^{-1}$ | Bact cells $mL^{-1}$ |
|---|---|---|---|---|---|---|
| Hard-bottom Winter (n=4) | 1 | 136 | 70.87±38.14% | 36.61±15.7% | 63.51±29.3% | 44.99±9.83% |
| | 2 | 155 | 52.84±26.68% | 42.68±16.61% | 66.56±17.28% | 57.83±5.36% |
| | 3 | 172 | 81.19±32.18% | 51.85±15.2% | 80.08±22.85% | 67.21±1.06% |
| | 4 | 184 | 59.98±14.83% | 65.93±21.96% | 84.98±29.53% | 72.18±15.44% |
| | 5 | 265 | 111.64±17.6% | 98.69±19.18% | 43.72±11.18% | 105.2±3.1% |
| | 6 | 396 | 144.31±14.54% | 101.49±10.85% | 70.49±8.69% | 109.81±15.44% |
| | 7 | 619 | 130.69±7.34% | 95.72±10.03% | 84.27±18.95% | 104.63±2.26% |
| | 8 | 896 | 100±0% | 100±0% | 100±0% | 100±0% |
| Hard-bottom Spring (n=2) | 1 | 114 | 35.3±9.99% | 117.36±41.5% | 58.89±17.43% | 47.36±13.32% |
| | 2 | 141 | 43.17±10.52% | 76.23±11.89% | 66.64±6.43% | 67.43±8.85% |
| | 3 | 164 | 57.96±7.32% | 96.87±13.58% | 101.02±40.64% | 73.83±6.01% |
| | 4 | 198 | 56.91±5.05% | 106.08±23.45% | 93.12±26.75% | 82.46±2.4% |
| | 5 | 247 | 88.35±4.1% | 141.72±25.79% | 309.41±226.39% | 98.7±12.99% |
| | 6 | 398 | 110.34±8.04% | 143.62±25.11% | 120.93±31.21% | 112.81±15.14% |
| | 7 | 627 | 96.31±5.94% | 143.57±35.3% | 103.03±19.03% | 93.4±4.83% |
| | 8 | 914 | 100±0% | 100±0% | 100±0% | 100±0% |
| Sand (n=2) | 1 | 107 | 153.45±7.99% | 195.92±10.09% | 231.64±61.72% | 132.79±0.02% |
| | 2 | 140 | 132.34±21.88% | 162.9±32.57% | 146.44±24.2% | 128.18±12.89% |
| | 3 | 164 | 126.72±33.22% | 146.53±30.42% | 137.76±23.38% | 125.84±11.98% |
| | 4 | 218 | 102.77±24.34% | 154.85±40.73% | 135.51±17.76% | 104.66±0.4% |
| | 5 | 294 | 96.72±20.68% | 171.36±81.74% | 114.45±5.22% | 106.72±19.56% |
| | 6 | 486 | 112.31±12.31% | 141.21±41.21% | 108.86±8.86% | 104.43±4.43% |
| | 7 | 627 | 120.14±0% | 161.44±0% | 106.4±0% | 106.13±0% |
| | 8 | 907 | 100±0% | 100±0% | 100±0% | 100±0% |
